# Efficient PCR-based gene targeting in isolates of the non-conventional yeast *Debaryomyces hansenii*

**DOI:** 10.1101/2023.07.11.548570

**Authors:** Sondos Alhajouj, Selva Turkolmez, Tarad Abalkhail, Zeena Hadi Obaid Alwan, D. James Gilmour, Phil J. Mitchell, Ewald H. Hettema

## Abstract

*Debaryomyces hansenii* is a yeast with considerable biotechnological potential as an osmotolerant, stress tolerant oleaginous microbe. However, targeted genome modification tools are limited and require a strain with auxotrophic markers. Gene targeting by homologous recombination has been reported to be inefficient but here we describe a set of reagents and a method that allows gene targeting at high efficiency in wild type isolates. It uses a simple PCR-based amplification that extends a completely heterologous selectable marker with 50 bp flanks identical to the target site in the genome. Transformants integrate the PCR product through homologous recombination at high frequency (>75%). We illustrate the potential of this method by disrupting genes at high efficiency and by expressing a heterologous protein from a safe chromosomal harbour site. These methods should stimulate and facilitate further analysis of *D. hansenii* strains and open the way to engineer strains for biotechnology.

## Introduction

The marine ascomycetes yeast *Debaryomyces hansenii* is a non-conventional oleaginous budding yeast. It is found in natural salty environments, and also on salted foods such as cheeses and cured meats, where it contributes to the development of flavour (Prista, Michan, Miranda, & Ramos, 2016). Its oleaginous physiology and its resistance to perchlorate to control growth of unwanted microorganisms in bioreactors are considered desirable traits for biotechnology. In addition, interest has grown within the scientific community to use it as a model to understand salt tolerance (Breuer & Harms, 2006; Prista et al., 2016). Most *D. hansenii* strains are haploid that can be diploid temporarily through autogamy (van der Walt, Taylor, & Liebenberg, 1977). Different isolates display heterogeneity in physiology and genome composition and in order to unlock and exploit this biological diversity for biotechnology, simple universal synthetic biology tools are essential for further development of *D. hansenii*.

Only a limited number of studies reported the development of research tools including a laboratory strain with auxotrophic markers, a replicating plasmid, gene disruption by homologous recombination using extensive flanking regions, random integration plasmids and CRISPR-Cas9 mediated gene targeting (Biswas, Datt, Aggarwal, & Mondal, 2013; Chawla, Kundu, Randhawa, & Mondal, 2017; Defosse et al., 2018; Minhas, Biswas, & Mondal, 2009; Minhas et al., 2012; Spasskaya et al., 2021; Strucko, Andersen, Mahler, Martinez, & Mortensen, 2021).

Homologous recombination was reported to be inefficient in *D. hansenii* (Minhas et al., 2009; Minhas et al., 2012; Spasskaya et al., 2021; Strucko et al., 2021) and long flanking regions were used for targeted integrations (Minhas et al., 2009; Minhas et al., 2012). This method relies on multiple plasmid construction steps to generate the gene targeting constructs and the auxotrophic markers *DhHIS4* or *DhARG1* (Biswas et al., 2013; Chawla et al., 2017; Minhas et al., 2009; Minhas et al., 2012) and therefore, this approach is mostly limited to the single *D. hansenii* strain DH9. One of the simplest methods of genomic modification is the PCR-mediated gene disruption method developed for gene targeting in *Saccharomyces cerevisiae* (Baudin, Ozier-Kalogeropoulos, Denouel, Lacroute, & Cullin, 1993). In this method, a selectable marker is amplified by PCR, extending the cassette with a 35-50 bp flanking region identical to the target site in the genome on either side. Upon transformation, these short flanking regions are sufficient to direct the selectable marker to the precise site in the genome by homologous recombination. This methodology opened the way for the systematic analysis of each gene and protein in *S. cerevisiae*. For instance, large arrays of gene deletion strains (for overview see (Giaever & Nislow, 2014) and genome wide libraries in which each ORF is tagged with, for instance GFP (Huh et al., 2003), mCherry (Yofe et al., 2016) or the TAP tag (Gavin et al., 2002) were generated and are available to the yeast research community. In a similar way, gene targeting in *Schizosaccharomyces pombe* and *Kluyveromyces lactis* have been developed (Kaur, Ingavale, & Bachhawat, 1997; Kooistra, Hooykaas, & Steensma, 2004). Initial studies in *S. cerevisiae* made use of auxotrophic markers. However, since these markers are identical to sequences in the host genome, gene conversions may restore the auxotrophy. Targeting efficiency was increased upon the use of heterologous selectable markers (Bahler et al., 1998; Wach, Brachat, Pohlmann, & Philippsen, 1994).

Here we describe an easy method for targeted gene deletion using a PCR mediated approach as previously developed for *S. cerevisiae*. Regions of 50 nt identity to the target site are sufficient to direct homologous recombination at high efficiency. We developed new selectable marker cassettes existing exclusively of heterologous DNA sequences that confer Hygromycin B or G418 resistance to *D. hansenii* transformants. Gene disruptions were obtained with high efficiency in 2 different isolates. Surprisingly, a third isolate showed no phenotypic effects of some of the gene disruptions and a wild type copy of the gene was also present, in addition to a disrupted gene copy. As low copy plasmids that segregate with high fidelity in *D. hansenii* are not available, we used a safe genome harbour site for the integrated expression of heterologous genes in this isolate. With these PCR based methods and reagents, genetic modification of *D. hansenii* isolates is now economic and easy to achieve and will pave the way to study this yeast into greater depth and develop it for biotechnological uses.

## Materials and Methods

### Strains, Growth Conditions and Media

*D. hansenii* NCYC102, NCYC3981 and NCYC3363 were obtained from Natural Collection of Yeast Cultures, Norwich Research Park, Norwich, UK (can be accessed via https://www.ncyc.co.uk/). Cells were grown in YM medium containing 0.3% w/v Yeast Extract (Formedium), 0.3% w/v Malt Extract (Oxoid), 0.5% w/v Peptone (Formedium) and 1% w/v D-Glucose (Fisher Scientific). For solid medium, 1.5% w/v Agar (Formedium) was added. Where antibiotic selection was required, hygromycin B (PhytoTech Labs) or G418 disulphate (Melford) were added into YM medium. The cells were grown at 25°C, either on solid media or in liquid cultures on a shaker at 200 rpm.

Yeast Minimal Arginine deficient (Arg-) or Adenine-deficient (Ade-) medium containing 0.5% w/v ammonium sulphate (BDH), 0.19% w/v Yeast Nitrogen Base without ammonium sulphate and amino acids (Sigma-Aldrich Y1251), 2% w/v D-Glucose, 2% w/v Agar for solid medium and 0.74 g/L Complete Supplement (CSM) DROP-OUT: -ARG or -ADE (Formedium).

For fluorescence microscopy, cells were grown to log phase in Yeast minimal complete medium containing 0.5% w/v ammonium sulphate (BDH), 0.19% w/v Yeast Nitrogen Base without ammonium sulphate and amino acids (Sigma-Aldrich Y1251), 2% w/v D-Glucose, 1% w/v Casamino Acids (Formedium), 20 μg/ml Uracil, 20 μg/ml Tryptophan and 30 μg/ml Leucine (Sigma-Aldrich).

*E. coli* DH5α cells were grown at 37°C, either as a liquid culture under shaking at 200 rpm or on solid media, using 2TY media which contains 1.6% w/v Tryptone (Formedium), 1% w/v Yeast Extract, 0.5% w/v NaCl (Fisher Scientific) and 2% w/v Agar for solid medium. Where required, Ampicillin Sodium Salt (MP Biomedicals) was added into the medium to the final concentration of 75 μg/ml.

### Molecular Biology techniques

Standard DNA manipulations were performed as described in Sambrook and Russell, 1982 except for where kits were used. Plasmids were constructed and amplified in *E*.*coli* DH5α. Plasmids were purified using QIAGEN Plasmid Miniprep kit, according to manufacturer recommendations. The different PCR polymerases and buffers were supplied by Meridian Bioscience (formerly Bioline). The oligonucleotides were supplied by Merck (formerly Sigma-Aldrich). The PCR protocols were performed as prescribed by the manufacturer and primers used in this research are described in Table 2. PCR products and plasmid digests were analysed by 0.7% agarose/TAE gel electrophoresis. Gels contained 0.5 μg/m Ethidium Bromide (Merck) to visualise DNA using a UV transilluminator supplied by GeneSys, and captured using GeneSys Software (Geneflow,UK). Molecular weight markers used were the 1 kb HyperLadder (from Bioline). T4 DNA ligase and buffer were purchased from New England Biolabs. The DNA sequences of newly made plasmids were confirmed by Sanger Sequencing, which was carried out by Source Bioscience. The samples were prepared and shipped to Source Bioscience Labs using their recommendations. Results were received as a Snapgene file and analysed by Clustal Omega database (Sievers et al., 2011) by aligning both estimated and obtained plasmid sequences.

### Generation of selectable cassettes

To generate the HygR marker plasmid, *Klebsiella pneumoniae* hygromycin B phosphotransferase ORF (*hph* ORF) was CTG codon adapted first, by changing all the CTG codons to other leucine codons. Codon-adapted *hph* ORF was placed in between *TEF1* promoter and terminator from *S. stipitis* (500 bp upstream and 250 bp downstream of *TEF1*, respectively). To generate the KanR cassette plasmid as a selectable marker, the bacterial kanamycin resistance (kanr) ORF from the *E. coli* transposon Tn903, which confers resistance to G418/Geneticin in eukaryotes, was CTG codon-adapted first and then placed under control of the *S. stipitis ACT1* promoter and terminator (500 bp upstream and 250 by downstream region of the *ACT1* ORF respectively). For both plasmids, the commonly used restriction sites within the expression cassette sequences were removed in a way that the corresponding amino acid sequences will not change. The selectable marker regions (promoter-antibiotics resistance ORF-terminator sequences) in both plasmids were flanked by multiple cloning sites to allow cloning, as well as loxP sites for future further recycling of the plasmids if necessary. Finally, both selectable markers were synthesised artificially and cloned into pUC19 by GenScript. For DNA sequences of various parts, see Table 1.

**Table 1.**
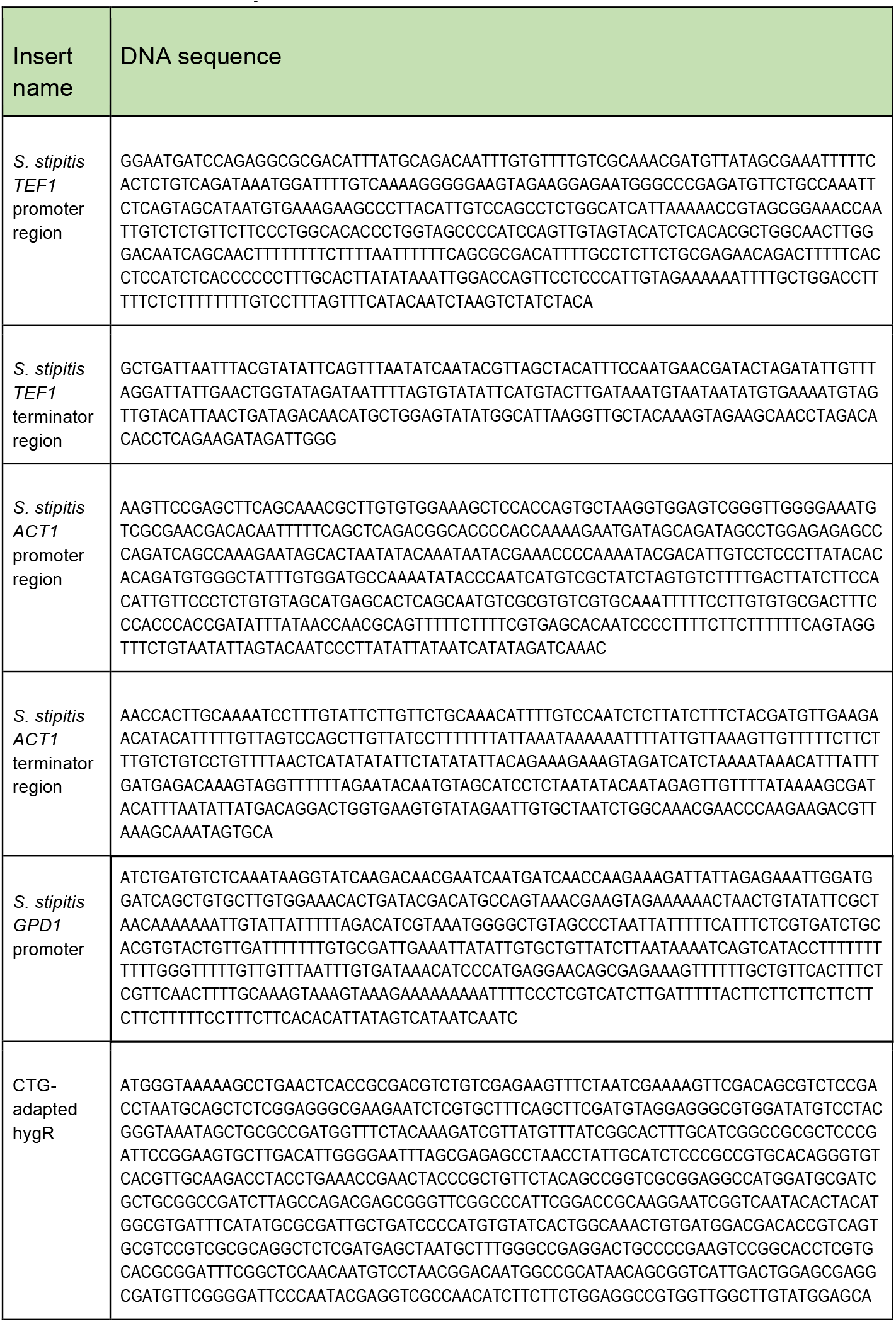
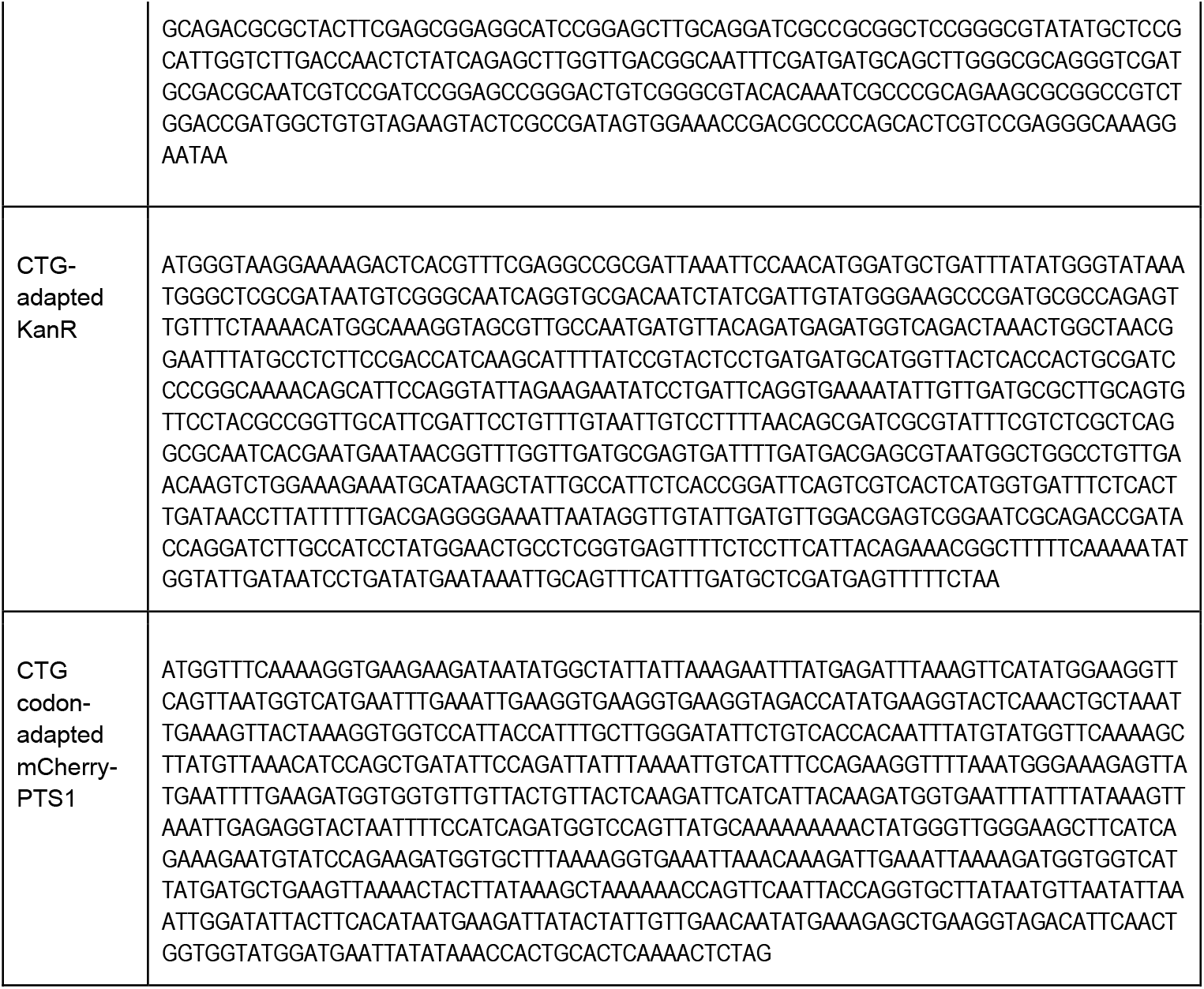
Promoter, terminator and antibiotics resistance ORF sequences that were used in this study.

**Table 2.**
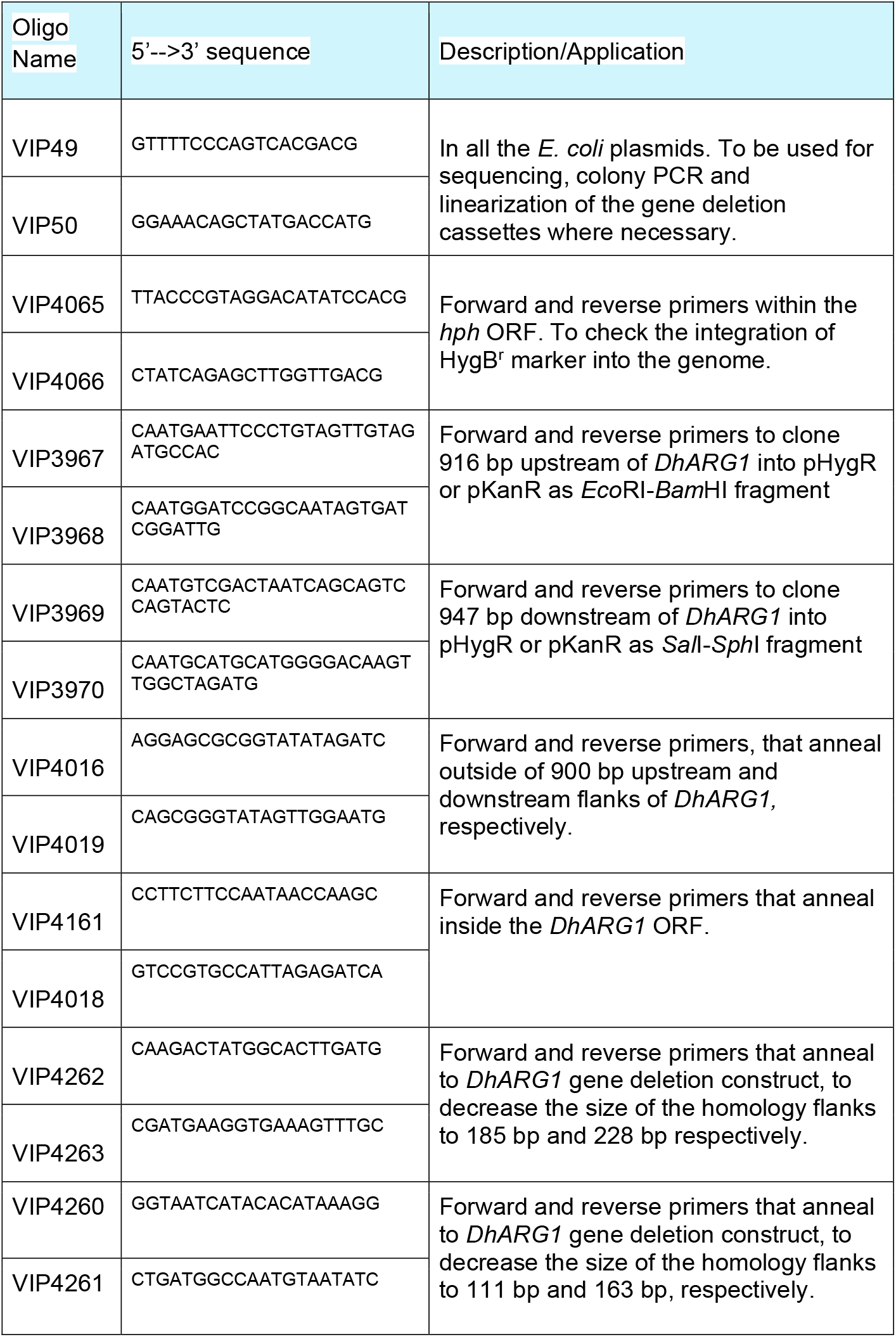
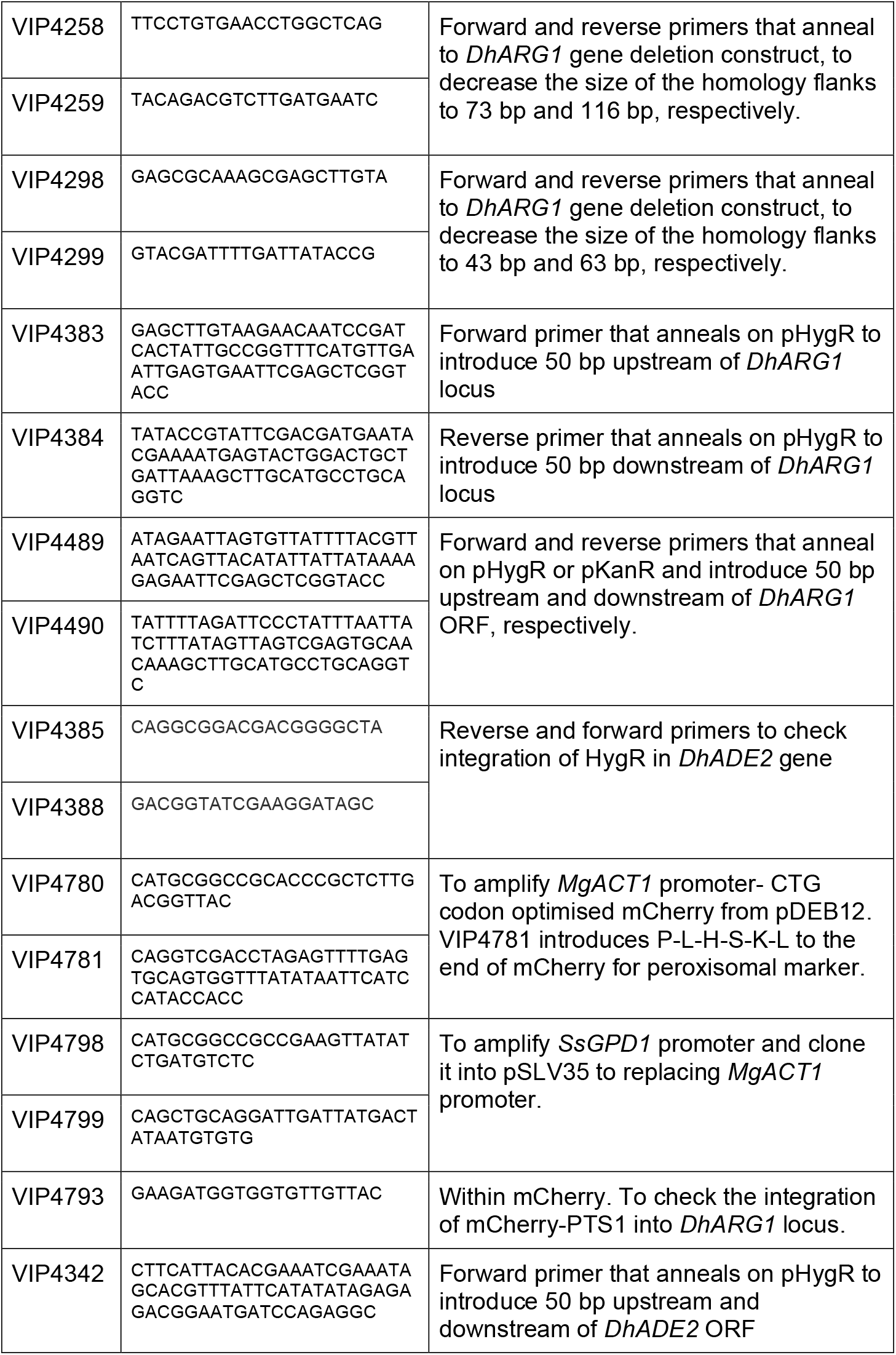
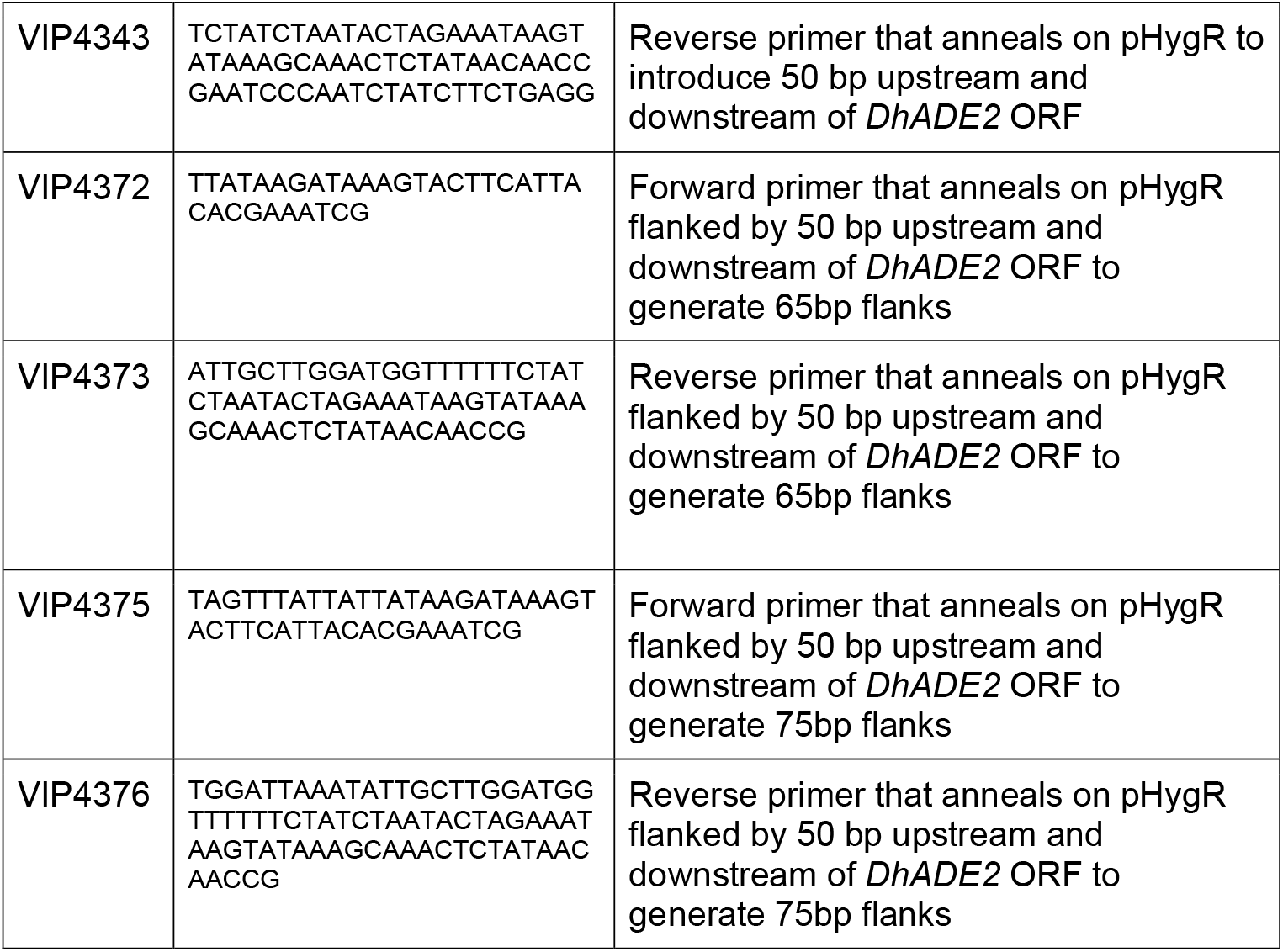
Oligonucleotides used in this study

**Table 3:** List of plasmids used in this study.

| Plasmid Name | Insert | Parental vector | Purpose | Source |
| --- | --- | --- | --- | --- |
| pHygR | loxP-HygR expression marker (*S. stipitis* *TEF1* promoter-CTG adapted hygR ORF-*S. stipitis* *TEF1* terminator)-loxP | pUC19 | To generate gene deletions/modifications in *D. hansenii* using HygR as a marker | GenScript |
| pKanR | loxP-KanR expression marker (*S. stipitis* *ACT1* promoter-CTG adapted KanR ORF-*S. stipitis* *ACT1* terminator)-loxP | pUC19 | To generate gene deletions/modifications in *D. hansenii* using KanR as a marker | GenScript |
| pSA4 | ~1 kb of upstream flank of *DhARG1* *EcoR1*-*BamH1* fragment | pKanR | Used as a backbone plasmid to generate peroxisomal marker plasmid with mCherry-PTS1. | This study |
|  | ~1 kb of downstream flank of *DhARG1* *SaI1*-*Sph1* fragment |  |  |  |
| pSA5 | ~0.9 kb of upstream flank of *DhARG1* *EcoR1*-*BamH1* fragment | pHygR | To generate *DhARG1* deletion cassette using long flanks of *DhARG1*. | This study |
|  | ~1 kb of downstream flank of *DhARG1* *SaI1*-*Sph1* fragment |  |  |  |
| pDEB12 | yemCherry from pAYCU257 (Defosse et al., 2018) *BamH1*-*SaI1* fragment | pAYCU272 (Defosse et al., 2018) | To swap the *StlacZ* in pAYCU272 with yemCherry of pAYCU257, after digesting both vectors with *BamH1* and *SaI1* | Defosse et al., 2018 |
| pSLV35 | *MgACT1* promoter-CTG codon adapted yemCherry-PTS1 *Not1*-*SaI1* fragment | pSA4 | To generate red peroxisomal marker construct plasmid backbone | This study |
| pSLV37 | *S. stipitis* *GPD1* promoter *Not1*-*Pst1* fragment | pSLV35 | To generate final peroxisomal marker construct behind *SsGPD1* promoter | This study |

### Plasmid construction

*ARG1* gene targeting constructs was generated by first amplifying the genomic region upstream of the *ARG1* gene using primers VIP3967 and 3968, which contain *Eco*RI and *Bam*HI restriction sites.The downstream region was amplified using VIP3969 and 3970, which contain *Sal*I and *Sph*I restriction sites, respectively. The PCR products were cloned into pHygR and pKanR using the restriction sites mentioned above. Peroxisomal marker plasmid was constructed as follows. The plasmids pAYCU257 and pAYCU272 were a gift from (Defosse et al., 2018). Both plasmids were double digested with *BamH*I and *Sal*I. Subsequently, the *Streptococcus thermophilus* lacZ in pAYCU272 was replaced with yemCherry from pAYCU257, which gave rise to pDEB12. Using pDEB12 as a template, the *Meyerozyma guilliermondii ACT1* promoter*-*CTG codon adapted yemCherry ORF sequence was amplified using primers VIP4780 and VIP4781, the latter adds the Peroxisomal Targeting Signal Type 1 (PTS1) -P-L-H-S-K-L to the 3’ end of the CTG codon-adapted mCherry. The resulting product was cloned into pSA4 (pKanR flanked by the long *DhARG1* homology arms) in between *Not*I and *Sal*I sites, which yielded pSLV35 plasmid. The *S. stipitis GPD1* promoter (500 bp upstream of *GPD1* ORF) was synthesised by Genscript and provided in pUC19. Using the oligonucleotides VIP4798 and VIP4799, the *S. stipitis GPD1* promoter was amplified by PCR to replace the *M. guilliermondii ACT1* promoter in pSLV35 by cloning, using *Not*I and *Pst*I sites. This yielded pSLV37. For plasmid design and DNA sequences of various parts see Table 1.

### Transformation of PCR products

Electroporation was based on a previously described method by (Minhas et al., 2009) with some small adaptations (Alwan, 2017). A fresh culture was diluted in 50 ml of YM medium at OD_600_ = 0.025-0.0125 and grown overnight. Next day, when cells reached OD_600_ =2.6-2.8, the cells were transferred into a sterile 50 ml screw cap tube and centrifuged for 5 min at 1700 g. The supernatant was discarded, and the cell pellet was resuspended in 1 ml of 50 mM sodium phosphate buffer, pH 7.5, containing 25 mM DTT. Cells were then incubated at 25°C for 15 min and centrifuged for 5 min at 1700 g. The cell pellet was resuspended in 8 ml sterile water (4°C) and centrifuged as previously. The supernatant was discarded, the cell pellet was resuspended in 1 ml sterile ice cold 1 M sorbitol and centrifuged as before. The supernatant was discarded, and the cells were resuspended in the remaining liquid to obtain a dense suspension (ca. 200 μl). Of the cell suspension, 40 μl was transferred to a microfuge tube and mixed with PCR product. Initial experiments used PCR product directly, but during this study it became clear that 500 ng EtOH-precipitated PCR product dissolved in 1-2 μl ultrapure water resulted in a consistently higher number of transformants. The mixture was placed in a precooled 2 mm electroporation cuvette (Geneflow,UK) and incubated on ice for 5 min. Electroporation was performed with an Equibio electroporator (a BioRad Gene Pulser) at 2.3 kV, 1000 Ω and 50 μF with pulse as exponential decay. Following electroporation, the cells were resuspended with 1 ml YM medium containing 0.1 M sorbitol. The cell suspension was transferred to a 2 ml microfuge tube. The samples were incubated for 4 hours at 25°C with shaking at 200 rpm. The cells were spread onto YM plates containing the appropriate antibiotic, and incubated at 25°C. After 3 days, single colonies were picked, and restreaked on appropriate selective media.

### Yeast total DNA isolation

Cells from a 3 ml overnight culture were harvested in a 2 ml screw cap tube (by centrifuging at 10625 rcf for 1 minute). The resultant pellets were washed with 1 ml sterile H_2_O. The pellets were resuspended in 200 μl TENTS solution (1% SDS w/v, 2% Triton X-100 v/v, 1 mM EDTA, 100 mM NaCl and 20 mM Tris/HCl (pH 8)), 200 μl phenol-chloroform-isoamyl-alcohol (pci, 25:24:1) and 200 μl 425-600 μm glass beads (BioSpec Products) were added into each tube. The tubes were placed into the bead beater (BioSpec Products) at maximum speed for 45 seconds, then centrifuged for 30 seconds at 10625 rcf. Another 200 μl TENTS solution was added into each tube followed by a brief vortex, the tubes were centrifuged at 10625 rcf for 5 minutes. The supernatants were transferred into Eppendorf tubes and another 200 μl phenol-chloroform-isoamylalcohol was added to each tube. Each tube was vortexed and centrifuged at 10625 rcf for 5 minutes. The supernatants (∼300 μl) were transferred to new tubes. Each supernatant was mixed with 1/10 volume 3 M NaAc (pH5.2) and 2.5X volume 100% EtOH, and the tubes were kept on ice for 15 minutes. Each tube was centrifuged at 4°C at 12000 rpm for 15 minutes, using the accuSpin Micro R centrifuge (Fisher Scientific). The supernatant was removed, and the pellets were washed with 300-500 μl 70% EtOH and centrifuged in Sigma 1-14 microfuge at maximum speed for 5 minutes. The supernatant was removed, and each pellet was resuspended with 200 μl 1X TE (at pH 7.4) + 2 μl RNAse (10 mg/ml), incubated at room temperature for 10 minutes. Then, they were mixed with 1/10 volume 3 M NaAc (pH 5.2) and 2.5X volume 100% EtOH and kept on ice for 15-30 minutes. Each tube was centrifuged at 4°C and 12000 rpm for 15 minutes, using the accuSpin Micro R centrifuge (Fisher Scientific). The pellets were washed with 300-500 μl 70% EtOH and centrifuged in Sigma 1-14 microfuge at maximum speed for 5 minutes. After the supernatants were discarded, the pellets were air dried. Finally, each pellet was resuspended in 50 μl TE (pH 7.4), and stored at -20°C.

#### Fluorescence microscopy

Live cell imaging of *D. hansenii* cells was performed as described previously for *S. cerevisiae* cells (Ekal, Alqahtani, & Hettema, 2023). Basically, cells were grown to log phase in minimal 2% glucose medium and analysed with a microscope (Axiovert 200M; Carl Zeiss) equipped with an Exfo X-cite 120 excitation light source, band pass filters (Carl Zeiss and Chroma Technology), a Plan-Apochromat 63×1.4 NA objective lens (Carl Zeiss) and a digital camera (Orca ER; Hamamatsu Photonics). Image acquisition was performed using Volocity software (PerkinElmer). Fluorescence images were collected as 0.5 μm *z*-stacks, merged into one plane using Openlab software (PerkinElmer) and processed further in Photoshop (Adobe). Bright-field images were collected in one plane and processed where necessary to highlight the circumference of the cells in blue.

## Results and discussion

### Development of a new dominant selectable marker cassettes for *D. hansenii*

We set out to improve the efficiency of HR mediated genome editing by generating a completely heterologous marker to allow for genome editing in wild type isolates of *D. hansenii* and to reduce the background of gene conversions by auxotrophic mutations. *D. hansenii* is a member of the CTG clade of yeasts, which also include *Candida albicans* and *Scheffersomyces stipitis*, wherein CUG codons, a common codon for leucine, is read as serine (Sugita & Nakase, 1999). We designed new selectable markers with the design of the hphMX and kanMX cassette for *Saccharomyces cerevisiae* gene targeting in mind (see Fig. 1A) (Goldstein & McCusker, 1999; Guldener, Heck, Fielder, Beinhauer, & Hegemann, 1996; Wach et al., 1994). The CTG codons of the *Klebsiella pneumoniae* hygromycin B phosphotransferase ORF, encoded by the *hph* gene (Gritz & Davies, 1983), were replaced by other leucine codons (for DNA sequence see Table 1). To allow the efficient expression of hph, a strong heterologous promoter (500 bp upstream region of the *S. stipitis TEF1* ORF) and terminator (250 bp downstream region of the *S. stipitis TEF1* ORF) sequences were selected. This marker cassette is flanked by *loxP* sites for future recycling. Additional restriction sites were introduced for convenience of cloning. The complete hygromycin B resistance cassette (HygR) was synthesised in pUC19 (Genscript) (Fig. 1A) and called pHygR. The KanR cassette was designed in a similar way. The bacterial kanamycin resistance (kanr) ORF from the *E. coli* transposon Tn903, which confers resistance to G418/Geneticin in eukaryotes, was placed under control of the *S. stipitis ACT1* promoter (500 bp upstream region of the *ACT1* ORF) and terminator (250 bp downstream of the *ACT1* ORF). The cassette is flanked by *loxP* sites that are surrounded by additional restriction sites. The CTG codons in the kanr ORF were replaced by CTA codons. The complete cassette (KanR) was synthesised in pUC19 (Genscript, for DNA sequence see Table 1) and called pKanR (Fig. 1A).

**Figure 1.**
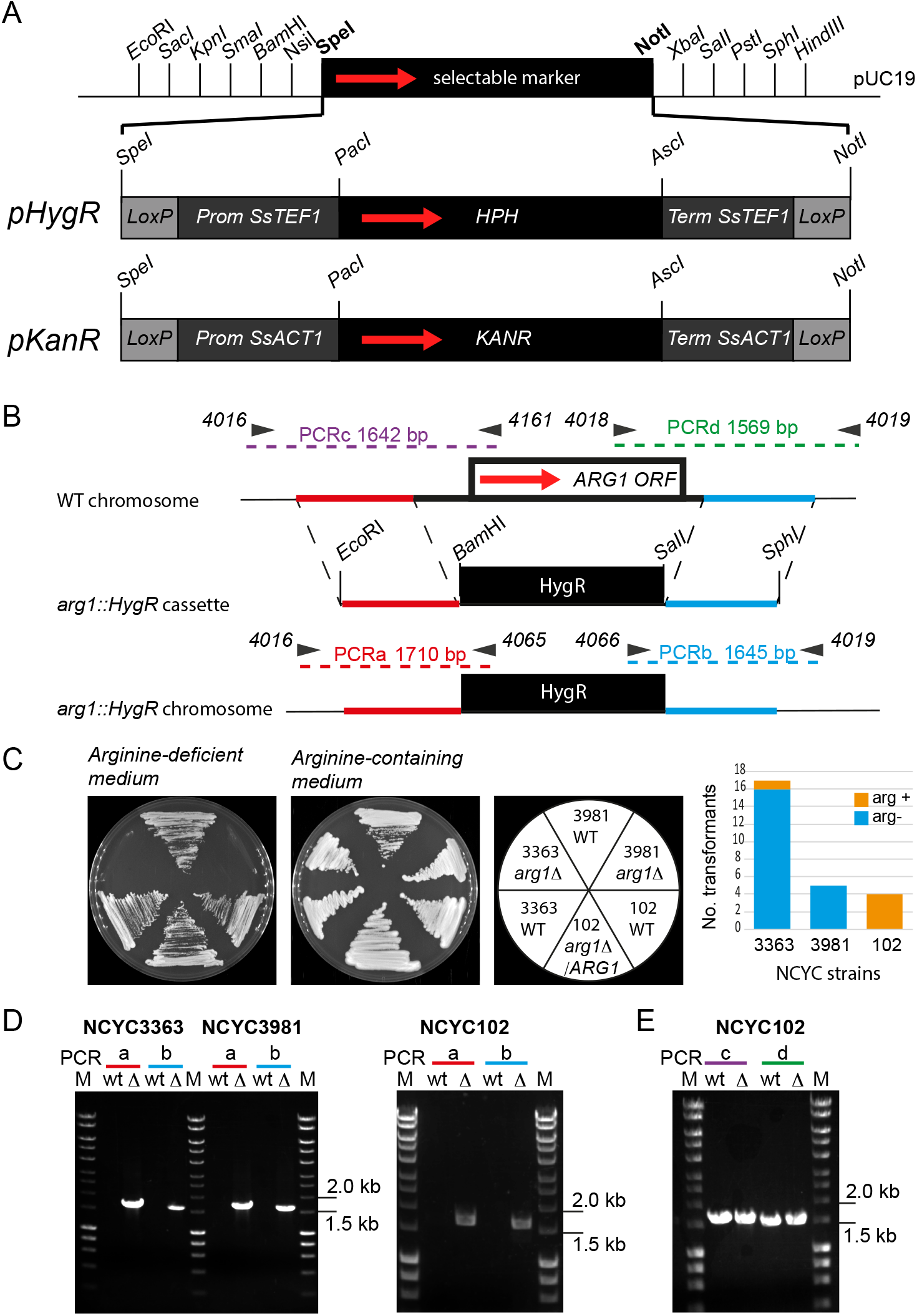
*ARG1* gene deletion strategy using completely heterologous antibiotic resistance marker cassettes. A) Schematic of antibiotic selection cassettes in pUC19 for use in *D. hansenii*, various elements not to scale. Restriction sites used for cloning homology arms and for identifying insert direction are indicated. Grey boxes represent the promoter (Prom) and terminator (Term) sequences of the *S. stipitis* translation elongation factor 1 gene or Actin 1 gene. Black boxes contain the *hph* gene from *Klebsiella pneumoniae* or the *kanr* gene from *E. coli*. pHygR and pKanR, plasmids encoding the new cassettes. B) *ARG1* gene deletion strategy using 0.9-1kb homology flanks. Black arrows containing numbers represent primers and the expected size of PCR products is indicated. Red and blue lines indicate flanking regions of the *ARG1* gene locus cloned into pHygR using restriction enzyme sites indicated. Red arrow, direction of transcription (A,B). C) Growth analysis of a selection of yeast strains and transformants as indicated on minimal arginine-deficient (arg-) and arginine-containing (arg+) medium. D,E) Agarose gel electrophoresis analysis of analytical PCR a,b,c and d as indicated in (B) on total DNA of wild type cells (wt) or hygromycin B resistant colonies (Δ). M, molecular weight marker.

To test the efficiency of the new HygR cassette in targeted gene deletions, we flanked the *hph* cassette with approximately 900 bp-1kb upstream and downstream regions of the arginine-succinate synthase encoding gene *ARG1* (Fig. 1B) and transformed this through electroporation into the three different natural *D. hansenii* isolates NCYC102, NCYC3363 and NCYC3981. Transformants were selected on 25 ug/ml Hygromycin B (for NCYC102) or 50 ug/ml Hygromycin B (for NCYC3981 and NCYC3363) and transformants were readily obtained. Hygromycin B resistant colonies were restreaked and subsequently tested for arginine auxotrophy. From the 5 and 17 NCYC3981 and NCYC3363 transformants obtained, 5 and 16 required arginine for growth, respectively (Fig. 1C). This suggested a high efficiency of targeted gene deletion in these two isolates. Correct integration was confirmed by analytical PCR (Fig. 1D). Surprisingly, none of the four NCYC102 transformants required arginine for growth (Fig. 1C) although the HygR cassette had integrated properly into the *ARG1* locus in all 4 of the colonies tested (one shown in Fig. 1D). Further PCR based analysis revealed the presence of an additional copy of *ARG1* in NCYC102 *arg1::HygR* transformants (Fig. 1E).

### Short homology arms direct efficient gene disruption

To test whether short homology arms promote homologous recombination in *D. hansenii*, oligonucleotides were designed to anneal on the original *ARG1* knock out cassette to achieve shorter flanks surrounding the *HygR* selection marker (see Fig. 2A). The differently sized *ARG1* deletion cassettes were amplified by PCR and were electroporated into NCYC3363. Transformants were selected on 50 ug/ml Hygromycin B. All PCR generated *ARG1* knock out cassettes yielded transformants with the majority of transformants requiring arginine for growth (Fig.2A). As homology arms of 43-63 bp still directed targeted deletion of the *ARG1* locus, we next designed oligonucleotides that directly annealed onto pHygR and contain 50 bp 5’ extensions identical to the DNA sequence of the *ARG1* locus (Fig. 2B,C). Using these oligonucleotides, the deletion cassette was amplified by PCR and was electroporated into NCYC3363 cells. We noticed variability between transformation efficiency of different PCR preparations and tested different PCR purification protocols. We found that electroporation of 500 ng of ethanol precipitated PCR product resulted in 17 to 25 transformants per transformation whereas electroporation of 500 ng PCR products cleaned up by a commercial PCR clean up kit resulted in a lower number of transformants (Fig 2D). Subsequently, we also transformed NCYC3981 using 500 ng ethanol precipitated PCR product and obtained between 10 and 21 transformants. (Fig. 2D). These conditions were used for further experiments (see also Materials and Methods for details). When we combine the results of these experiments, 106 of the 145 NCYC3363 transformants (73%) and 38 of the 47 NCYC3981 transformants (80%) failed to grow on arginine-deficient medium. We confirmed correct integration of the *HygR* cassette into the *ARG1* locus in two randomly selected arginine auxotrophs of each strain background (Fig. 2E). We conclude that PCR mediated gene disruption is a very efficient way of generating mutants in *D. hansenii* NCYC3363 and NCYC3891 cells.

**Figure 2.**
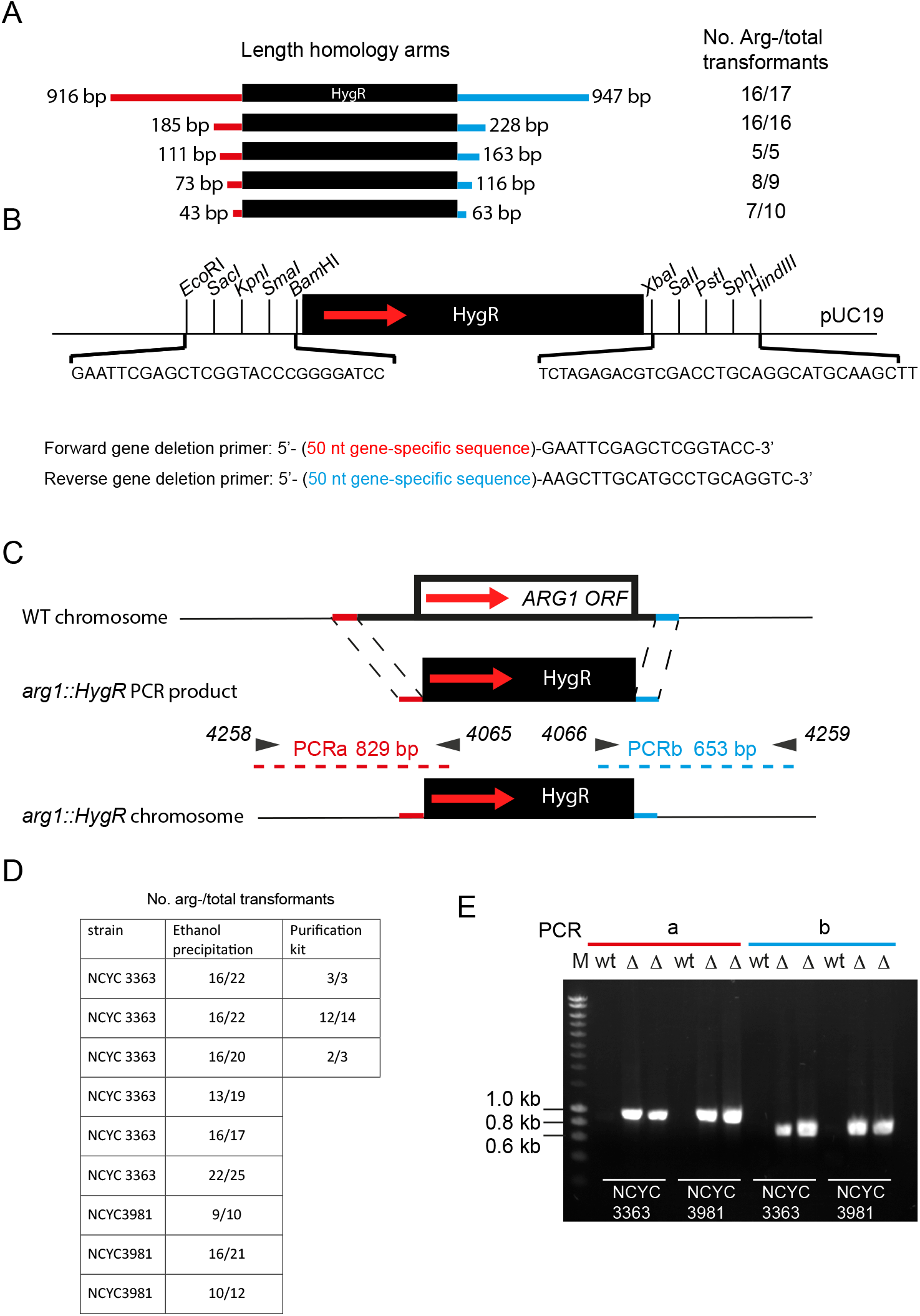
Short homology arms direct gene targeting with high efficiency. A) Schematic showing the reduction of homology arm length of the *ARG1* knock out construct and the resulting number of arginine auxotrophs (Arg-) among the total hygromycin B resistant transformants in the strain NCYC3363. B) Schematic showing design of primers to amplify the completely heterologous antibiotic resistance cassettes for gene deletions using 50 bp homology arms. C) *ARG1* gene deletion strategy using 50 bp homology flanks generated by direct PCR on pHygR. Red arrow, direction of transcription. Black arrows containing numbers represent primers and the expected size of PCR products is indicated. Red and blue lines indicate 50 bp flanking regions just upstream and downstream of the *ARG1* ORF.D) Table of transformant frequency and arginine auxotrophy during tests of PCR product purification. E) Agarose gel electrophoresis analysis of PCR a and b products as indicated in (C) on total DNA of wild type cells (wt) or two hygromycin B resistant colonies (Δ) in the strains NCYC3363 and NCYC3891. M, molecular weight marker.

To test whether PCR based gene deletion was also efficient at other loci, we tested the *ADE2* locus. Three different *ADE2* KO cassettes were produced through PCR on p*HygR*. The cassettes varied in the length of homology arms (75 bp, 65 bp and 50 bp) identical to the regions directly upstream and downstream of the *ADE2* ORF (Fig. 3A). The PCR products were ethanol precipitated and 500 ng electroporated into NCYC3363 and NCYC3981 cells, and transformants were selected on hygromycin B-containing media. Colonies were then streaked onto Hygromycin B -containing medium and grown for 2 days before restreaking onto adenine-deficient (ade-) plates (Fig. 3B). The vast majority of transformants (>90%) in NCYC3363 and NCYC3981 failed to grow on ade-medium (Fig. 3B). Adenine auxotrophs accumulate a brown,reddish intermediate in the adenine biosynthetic pathway, hence the colour difference between adenine auxotrophs and prototrophs (Fig. 3B). PCR analysis confirmed correct integration of the *HygR* cassette into the *ADE2* locus in a randomly selected adenine auxotroph of each strain background (Fig. 3C). We conclude that gene targeting occurs efficiently at both the *ARG1* and the *ADE2* locus using short regions of homology.

**Figure 3.**
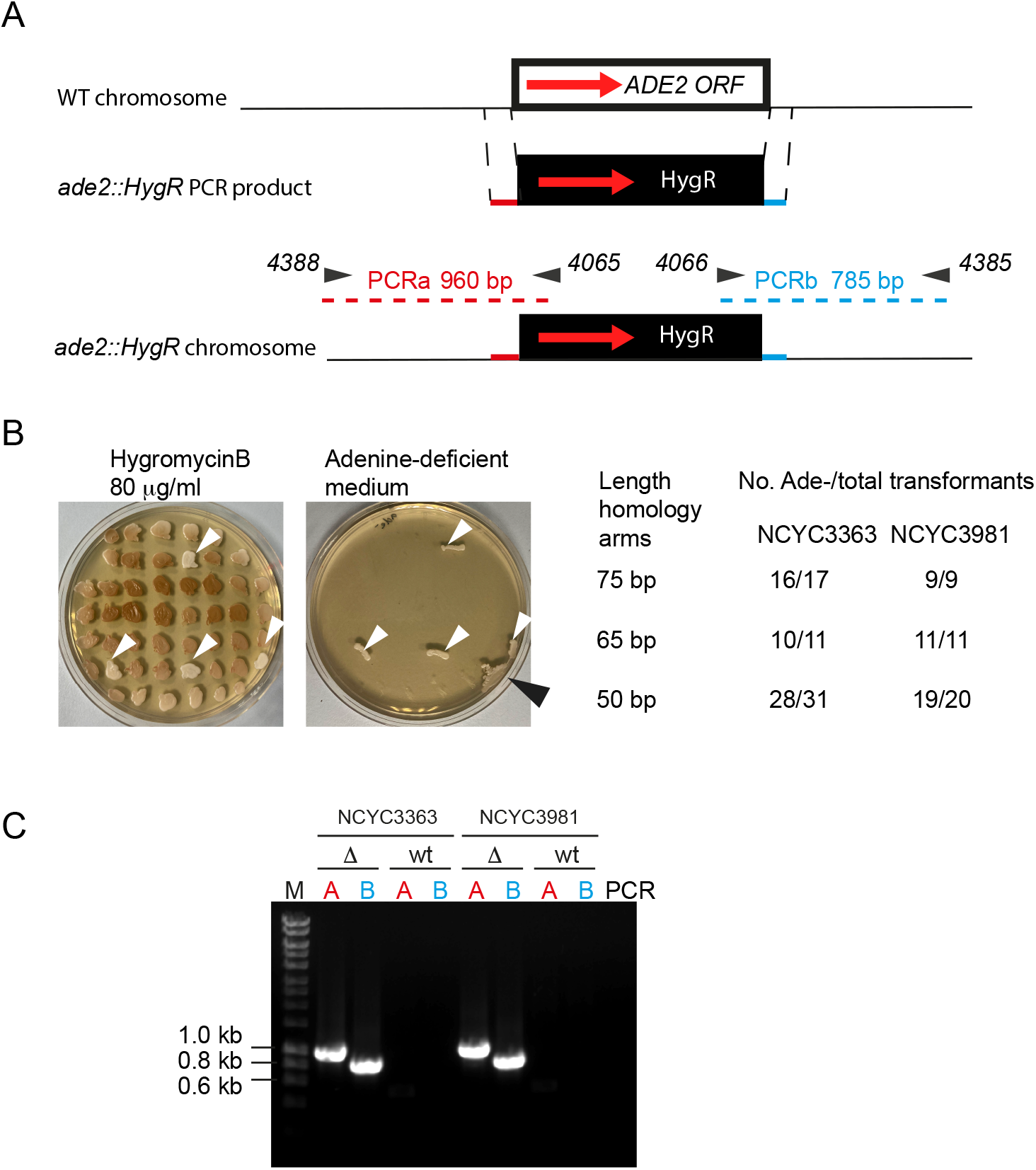
*ADE2* gene deletion using 50 bp, 65bp and 75 bp homology flanks generated by direct PCR on pHygR. Red arrow, direction of transcription. Black arrows containing numbers represent primers and the expected size of PCR products is indicated. Red and blue lines indicate 50 bp flanking regions just upstream and downstream of the *ADE2* ORF. B) Growth analysis of a collection of 49 NCYC3361 transformants patched onto hygromycin B-containing medium, grown for 2 days at 25°C and from there they were patched on minimal adenine-deficient (ade-) medium and incubated for another 2 days at 25°C. Note the white patches on the hygromycin B medium, correspond to adenine prototrophs as indicated by white arrow heads. Black arrowhead indicates a patch of untransformed wt cells added as positive control onto ade-plates, as these cells do not grow on Hygromycin B plate. The resulting number of adenine auxotrophs (Ade-) among the total hygromycin B resistant transformants in the strain NCYC3363 NCYC3981 is indicated. C) Agarose gel electrophoresis analysis of PCR a and b products as indicated in (A) on total DNA of wild type cells (wt) or hygromycin B resistant colonies (Δ) in the strains NCYC3363 and NCYC3891. M, molecular weight marker.

### Safe harbour site for heterologous expression in the *ARG1* locus

As low copy plasmids that segregate with high fidelity in *D. hansenii* are not available, and since random integration of expression cassettes in the genome may affect normal cell function and result in various levels of expression, we explored the expression of heterologous proteins by insertion of expression cassettes into one of the *ARG1* locus of NCYC102. As a test we generated a universal peroxisomal marker by appending the red fluorescent protein mCherry with a peroxisomal targeting signal type 1 (PTS1). Peroxisomes are small organelles that post-translationally import proteins into their lumen and that are gaining interest in synthetic biology circles to house new biosynthetic pathways for metabolic engineering purposes (see for instance Cross et al., 2017; DeLoache, Russ, & Dueber, 2016; Dusseaux, Wajn, Liu, Ignea, & Kampranis, 2020; Naseri, 2023). Using the *KanR* cassette flanked by long *ARG1* homology arms (Fig. 4A) as a starting point, we subsequently inserted mCherry-PTS1 under control of the *S. stipitis GPD1* promoter into this cassette. The cassette was PCR amplified and transformed into NCYC102 cells. G418 resistant colonies were selected, restreaked and grown into liquid media before imaging. As expected, a typical punctate fluorescent pattern was observed in the transformants (Fig. 4B). Proper integration into the *ARG1* locus was confirmed by PCR (Fig. 4C). We conclude the *ARG1* locus in NCYC102 cells provides a safe genome landing site for the expression of heterologous genes. And as NCYC102 has multiple copies of *ARG1*, the cells do not become arginine auxotrophs.

**Figure 4.**
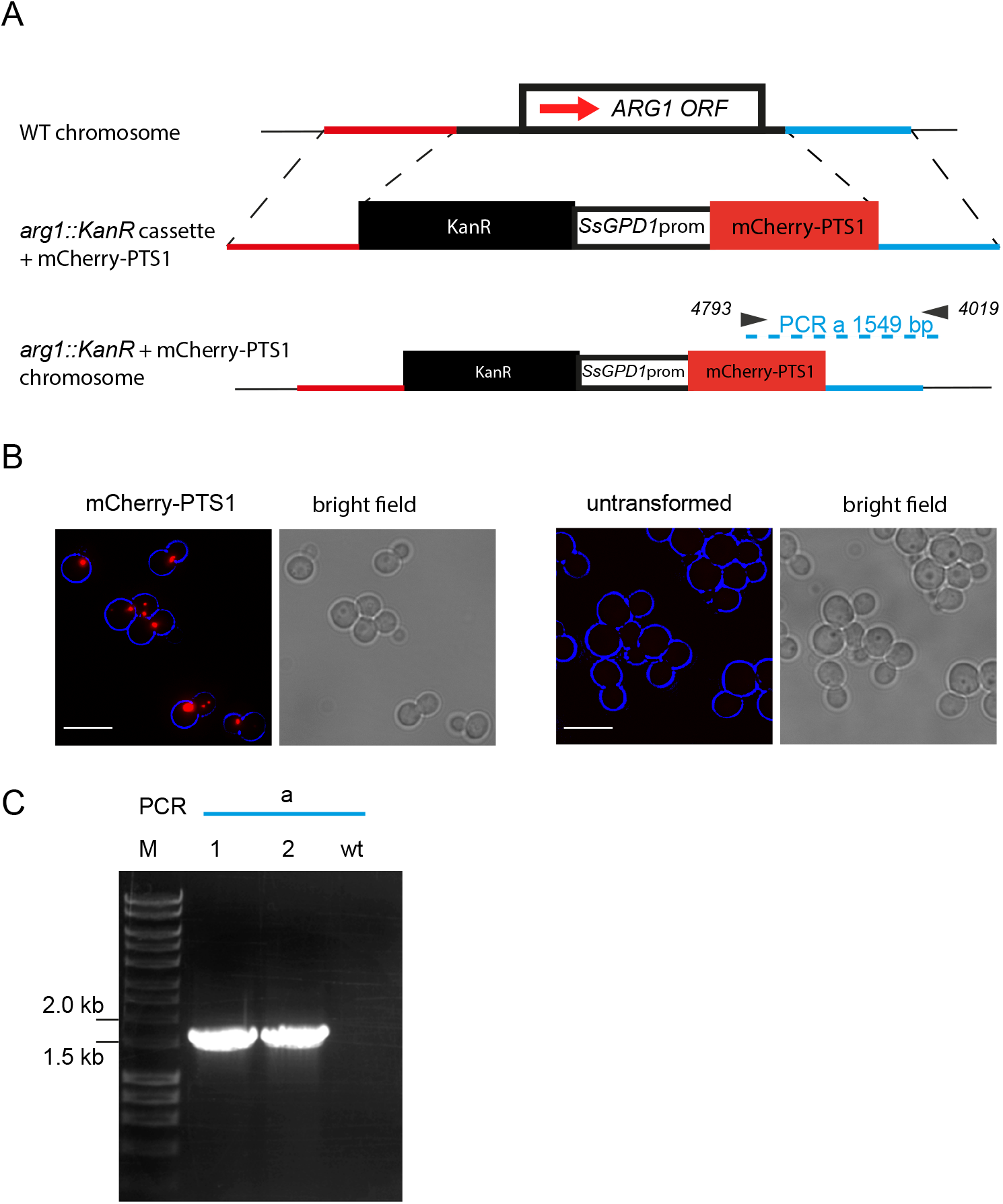
Strategy for heterologous expression through integration into the *ARG1* locus A) Schematic of the expression array of the KanR cassette followed by the mCherry-PTS1 expression unit, flanked by 0.9-1kb homology regions to the *ARG1* locus to stimulate integration through homologous recombination. The CTG codon adapted mCherry ORF was extended with a C-terminal peroxisomal targeting signal type I (PTS1) and expression is controlled by the *S. stipitis GPD1* promoter region (Prom). Red arrow, direction of transcription. Black arrows containing numbers represent primers and the expected size of PCR products is indicated. Red and blue line indicated upstream and downstream flanking regions of the *ARG1* gene locus, respectively. B) Epifluorescence microscopy image of NCYC102 cells transformed with mCherry-PTS1 that were grown for extended periods in log phase. Image is a flattened *z*-stack. Cell circumference is labelled in blue. Scale bar: 5 μm. C) Agarose gel electrophoresis analysis of PCR product as indicated in (A) on total DNA of wild type cells (wt) or two G418-resistant colonies (1, 2) that are showing punctate mCherry labelling. M, molecular weight marker.

## Conclusion

Here we describe an easy method for efficient gene targeting in *D. hansenii* isolates through homologous recombination using short flanks of only 50 bp. Although previous studies suggested that gene targeting through homologous recombination requires long regions of homology to target sites and occurs at low efficiency, we found this was not the case in our experiments as *ARG1* deletions in NCYC3363 and NCYC3981 together resulted in only 1 out of 22 transformants to be mistargeted. One likely explanation for this is that in previous studies auxotrophic markers were used. Since these markers are identical to the DNA sequences in the host genome, gene conversions may restore the auxotrophy, thereby reducing targeting efficiency as has previously been observed in *S. cerevisiae* (Wach et al., 1994). The high efficiency of targetted gene disruption prompted us to test whether short homology arms could stimulate gene targeting and indeed we found it could. However, we noticed a drop in the number of transformants but through optimisation of the electroporation protocol, we now routinely obtain between 15-50 per electroporation using 500 ng PCR product. In line with our findings, double strand breaks (DSB) induced by CRISPR-Cas9 could be repaired by homologous recombination using 90 bp oligonucleotides in *D. hansenii* (Spasskaya et al., 2021), further illustrating that short homologous sequences are sufficient to mediate accurate homologous recombination in this yeast. One surprising observation was that the strain NCYC102 appears to have multiple copies of the *ARG1* gene. We use this observation to our advantage to generate a safe site for expression of heterologous genes. *D. hansenii* isolates are assumed to be haploid, but variations in genome size have been reported and some strains are diploid or even aneuploid (Jacques et al., 2010; Link, Lulf, Parr, Hilgarth, & Ehrmann, 2022; Petersen & Jespersen, 2004). We have found several additional genes in this strain to still contain a WT copy after disruption of one copy, including *ADE2* and *FOX2* (not shown). This suggests that NCYC102 may be diploid or aneuploid. The use of PCR-based genome modifications is a very powerful tool for systematic approaches in the analysis of *D. hansenii* and the use of heterologous selectable markers allows to study not only the laboratory strain DH9 but also wild isolates.

## Acknowledgements

A PhD scholarship was awarded to S. Alhajouj by the Royal Embassy of Saudi Arabia Cultural Bureau in London and Jouf University Saudi Arabia. A PhD scholarship was awarded to T. Abalkhail by The Royal Embassy of Saudi Arabia Cultural Bureau in London and King Saud University Saudi Arabia. A PhD scholarship was awarded to Z.H.O. Alwan by the Iraqi Ministry of Higher Education and Scientific Research.

## Notes

### Competing Interest Statement

The authors have declared no competing interest.

